# Glutamatergic synaptic deficits in the prefrontal cortex of the Ts65Dn mouse model for Down syndrome

**DOI:** 10.1101/2023.02.22.529415

**Authors:** Aurore Thomazeau, Olivier Lassalle, Olivier J. Manzoni

**Author notes:** Correspondence, Address for editorial correspondence: Aurore Thomazeau, IPMC UMR7275, 660, route des Lucioles, Sophia Antipolis, 06560 Valbonne, France.

## Abstract

Down syndrome (DS), the most common form of intellectual disability, is a chromosomal disorder caused by having all or part of an extra chromosome 21, leading to intellectual disability. Contrary to the extensive research on the Ts65Dn mouse model of DS in the hippocampus, the synaptic foundation of prefrontal cortex (PFC) malfunction in individuals with DS, including working memory deficits, remains largely unclear. A previous study on mBACtgDyrk1a mice, which overexpress the *Dyrk1a* gene, showed that this overexpression negatively impacts spine density and synaptic molecular composition, causing synaptic plasticity deficits in the PFC. By comparing Ts65Dn mice, which overexpress multiple genes including *Dyrk1a*, and mBACtgDyrk1a mice, we aimed to better understand the role of different genes in DS. Results from electrophysiological experiments (i.e., patch-clamp and extracellular field potential recordings ex vivo) in Ts65Dn PFC male mice revealed modifications of intrinsic properties in layer V/VI pyramidal neurons and the synaptic plasticity range. Thus, long-term depression was abolished in Ts65Dn, while synaptic or pharmacological long-term potentiation were fully expressed in Ts65Dn mice. These results, illustrating the phenotypic divergence between the polygenic Ts65Dn model and the monogenic mBACtgDyrk1a model of DS, highlight the complexity of the pathophysiological mechanisms responsible for the neurocognitive symptoms of DS.

## 1. Introduction

Down syndrome (DS), the most common form of intellectual disability, is a chromosomal disorder resulting from the presence of all or part of an extra chromosome 21. The presence of an additional copy of the ∼ 300 normal genes present on chromosome 21 leads to multiple significant clinical phenotypes. The common feature among all individuals with DS is intellectual disability, reflected by characteristic and specific deficits in learning and memory (Chapman and Hesketh, 2000). The neural mechanisms underlying these alterations are still poorly understood, and may include defects in neural network formation, information processing and brain plasticity.

Mouse models of neurological disorders hold great promise in helping to identify the underlying cellular mechanisms of these conditions. To this end, several animal models that resemble the alterations found in DS have been created. The most well-researched mouse model for DS is the Ts65Dn mouse, which has a segmental trisomy of mouse chromosome 16 and carries three copies of genes that are equivalent to those found on human chromosome 21 (Hsa21), many of which are conserved between mice and humans (Davisson et al., 1990;Gardiner et al., 2003;Sturgeon and Gardiner, 2011;Rueda et al., 2012). The trisomic region includes ∼140 genes and overlaps largely with the region of Hsa 21 considered to be responsible for many DS phenotypes, including intellectual disability (Korenberg et al., 1994). Indeed, the Ts65Dn mice exhibits behavioural, cellular, and molecular phenotypes relevant for DS, such as alterations in working memory, deficits in long-term memory, hyperactivity, disrupted neurogenesis and synaptic plasticity, and a general over-inhibition (Holtzman et al., 1996;Gardiner and Davisson, 2000;Belichenko et al., 2004;Kleschevnikov et al., 2004;Belichenko et al., 2007;Rueda et al., 2012;Ruiz-Mejias, 2019). Most studies on synaptic plasticity have focused on the hippocampus, a brain structure with a major role in learning and memory (Cramer and Galdzicki, 2012).

The prefrontal cortex (PFC) is widely recognized as the area of the brain responsible for the control and coordination of executive cognitive processes such as planning, cognitive flexibility, working memory, and emotional behaviour (Goldman-Rakic, 1990;Seamans et al., 1995). PFC malfunction is a prevalent aspect of several neuropsychiatric diseases, including DS (Goto et al., 2010). Neuropsychological evaluations of individuals with DS have shown deficits in executive functions (Lanfranchi et al., 2010). The precise role of the triplication of these approximately 140 genes in PFC malfunction and the synaptic basis for the working memory impairments seen in individuals with DS is still not fully understood (Pennington et al., 2003;Lanfranchi et al., 2010).

In this study, the intrinsic characteristics and glutamatergic synaptic plasticity of deep-layer pyramidal neurons in the PFC of adult male Ts65Dn mice, were analysed. The focus was on male Ts65Dn mice only, as significant over-expression of DYRK1A in the cerebral cortex has been detected in male mice, but not in females (Hawley et al., 2022). The findings revealed modifications in the intrinsic properties of pyramidal neurons and specific changes in synaptic plasticity. Our results uncover functional abnormalities in the PFC of male Ts65Dn mice that may play a role in the multiple cognitive and behavioural impairments seen in individuals with DS.

## 2. Materials and methods

### 2.1. Animals and Ethics Statement

All animal experiments were performed according to the criteria of the European Communities Council Directive (86/609/EEC). The B6EiC3Sn.BLiA-Ts(1716)65Dn/DnJ [known as Ts65Dn] mice were obtained from Dr. Jean Delabar’s laboratory. Ts65Dn mice (Davisson et al., 1993) were maintained on a B6/C3H background and genotyped as described previously (Reinholdt et al., 2011). All mice were weaned at 21 days. After weaning, they were caged socially in same-sex groups. Male Ts65Dn mice and wild type littermate controls were used at 4 to 6 months of age.

### 2.2. Acute prefrontal cortex slice preparation

PFC slices were prepared as described (Lafourcade et al., 2011). Mice were anesthetized with isoflurane and decapitated. The brain was sliced (300 μm) in the coronal plane (Integraslice, Campden Instruments, Leicester, U.K.) and maintained in physiological saline (4°C). Slices were stored for 30 min at 32–35°C in artificial cerebrospinal fluid (ACSF) containing 126 mM NaCl, 2.5 mM KCl, 2.4 mM MgCl2, 1.2 mM CaCl2, 18 mM NaHCO3, 1.2 mM NaH2PO4 and 11 mM glucose, equilibrated with 95% O2/5% CO2. Slices were stored at 22 ± 2 °C until recording.

### 2.3. Electrophysiology

Whole-cell patch-clamp and field excitatory postsynaptic potential (fEPSP) were recorded from layer V/VI pyramidal cells in coronal slices of mouse prelimbic PFC (Lafourcade et al., 2011;Thomazeau et al., 2014). For recording, slices were superfused (2 ml/min) with ACSF at 32–35 °C. Picrotoxin (100 μM) was added to block GABA-A receptors. To evoke synaptic currents, 150-200 μs stimuli were delivered at 0.1 Hz through a glass electrode placed in layer II/III. The glutamatergic nature of the fEPSP was confirmed at the end of the experiments using the glutamate receptor antagonist 6,7-dinitroquinoxaline-2, 3-dione (DNQX, 20 μM), that blocked the synaptic component without altering the non-synaptic component (not shown). LTD was induced by 10 min stimulation at 10 Hz (Lafourcade et al., 2011). The presynaptic adenylate cyclase (AC)/cAMP signaling pathway-dependent facilitation was induced by bath application of the AC activator forskolin at 10 μM. NMDAR-LTP was induced using a theta-burst stimulation (TBS) protocol consisting of five trains of burst with four pulses at 100 Hz, at 200 ms interval, repeated four times at intervals of 10 s (Thomazeau et al., 2014). For whole-cell patch-clamp, pyramidal neurons were visualized using an infrared microscope (BX-50, Olympus). Experiments were performed with electrodes containing 128 mM potassium gluconate (KGlu), 20 mM NaCl, 1 mM MgCl2, 1 mM EGTA, 0.3 mM CaCl2, 2 mM Na2+-ATP, 0.3 mM Na+-GTP, 10 mM glucose buffered with 10 mM HEPES, pH 7.3, osmolarity 290 mOsm. Electrode resistance was 4-6 MOhm. If access resistance (no compensation, <25 MOhm) changed by >20%, the experiment was rejected. To perform current–voltage (I–V) curves and to test neuronal pyramidal neuron excitability, a series of hyperpolarizing and depolarizing current steps were applied immediately after breaking in the cell.

### 2.4. Data acquisition and analysis

Due to the strong phenotypes, the experimenters were not blind to the genotypes, but no data were excluded before statistical analysis (GraphPad Software Inc., La Jolla, CA). Data were recorded on a MultiClamp700B, filtered at 2 kHz, digitized (20 kHz, DigiData 1440A, collected using Clampex 10.2 and analysed using Clampfit 10.2 (all from Molecular Device, Sunnyvale, USA). Both area and amplitude of fEPSPs and EPSCs were computed. The magnitude of LTD and LTP was calculated respectively 30-35 min and 35-40 min after induction as percentage of baseline responses. To perform current–voltage (I– V) curves and to test the excitability of MSN, a series of hyperpolarizing and depolarizing current steps were applied immediately after 10 breaking in the cell. Membrane resistance was estimated from the I–V curve around resting membrane potential as previously described (Kasanetz and Manzoni, 2009;Thomazeau et al., 2014).

### 2.5. Drugs

Drugs were added at the final concentration to the ACSF. Picrotoxin was from Sigma (St. Quentin Fallavier, France). DNQX was from the National Institute of Mental Health’s Chemical Synthesis and Drug Supply Program (Rockville, MD, USA). Forskolin was from Tocris (Bristol, UK).

### 2.6. Statistical analysis

The value n corresponds to the number of individual animals in the electrophysiology experiments. All values are given as mean ± s.e.m. Statistical analysis was performed with Prism 7.0 (GraphPad Software Inc., La Jolla, CA). Two sample comparisons were made with the Mann-Whitney t test. Statistical significance was set at P < 0.05.

## 3. Results

### 3.1. Intrinsic properties of PFC pyramidal neurons are changed in Ts65Dn mice

Intrinsic properties of pyramidal neurons in layers V/VI of the cerebral cortex were recorded in the “current-clamp” mode. The curves showing the relationship between the current injected into the neuron and the change in membrane potential revealed a difference between Ts65Dn mice and wild-type euploid littermates. Thus, pyramidal neurons of aneuploid mice showed an augmented inward rectification (**Figure 1A**). However, no change in membrane potential was found (**Figure 1B**). The rheobase (i.e., the minimum current intensity required to induce a first action potential) was higher in Ts65Dn mice, reflecting a decrease in membrane excitability (**Figure 1C**). Finally, the number of action potentials emitted following incremental current injection was similar in the two mouse strains (**Figure 1D**).

**Figure 1.**
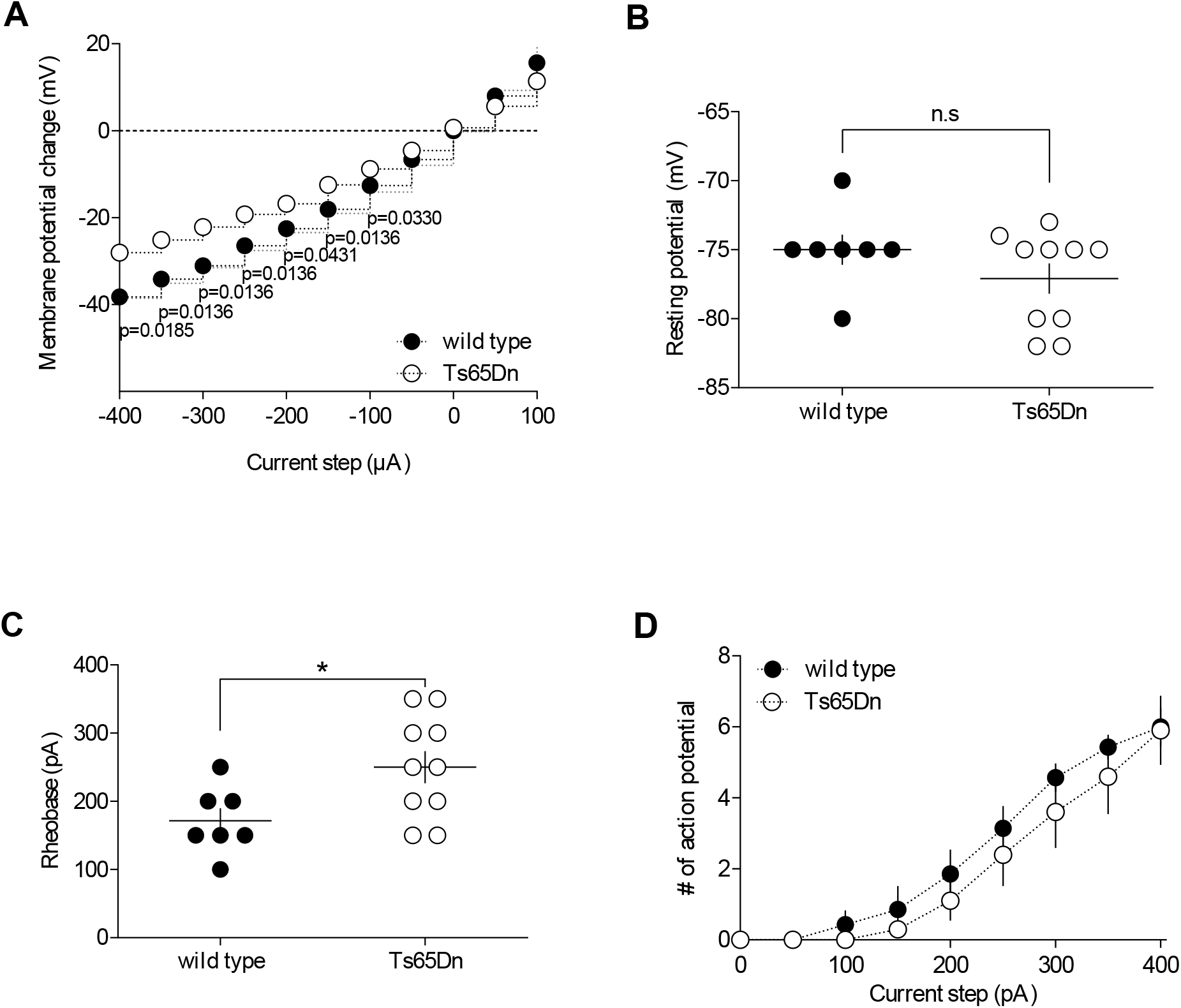
The intrinsic properties of pyramidal neurons in layers V/VI of the cerebral cortex are altered in Ts65Dn mice. **A**. Summary of current-voltage (I-V) curves recorded in pyramidal neurons of both strains showing an increase in inward rectification in Ts65Dn mice (white symbols; n = 10) compared to wild-type mice (black symbols; n = 7). Resting membrane potential is similar in both genotypes (**B**. -75.00 ± 1.09, wild-type mice, black symbols; -77.10 ± 1.10, Ts65Dn mice, white symbols; P > 0.05, Mann-Whitney test). In contrast the rheobase was higher in Ts65Dn mice (**C**. 171.40 ± 18.44, wild-type mice, black symbols; 250.00 ± 23.57, Ts65Dn mice, white symbols; P = 0.0438, Mann-Whitney test). The horizontal line represents the mean value. **D**. The summary of current-discharge curves indicates that the number of action potentials elicited in response to current injection steps is similar in pyramidal neurons of both strains. Error bars represent standard error to the mean.

### 3.2. Synaptic plasticity is partially affected in the Ts65Dn mouse model

Long-term changes of synaptic efficacy are thought to play pivotal roles in cognition, learning, and memory formation. We next compared in our mouse groups, opposite forms of synaptic plasticity that rely on distinct mechanisms, namely endocannabinoid-mediated long-term depression (eCB-LTD), presynaptic adenylyl cyclase/cyclic adenosine monophosphate signaling pathway-dependent facilitation (AC/cAMP-facilitation), and N-methyl-D-aspartate receptor-dependent long-term potentiation (NMDAR-LTP).

#### 3.2.1. Failed induction of endocannabinoid-dependent long-term depression in Ts65Dn mice

The eCB system is a major player in synaptic plasticity that may participate in the etiology of PFC-dependent mood disorders (Hill et al., 2009;Scheyer et al., 2017). In response to neuronal activity, eCBs are released by the postsynaptic compartments and retrogradely activate presynaptic cannabinoid receptors to induce long-term depression (LTD) (Robbe et al., 2002;Puente et al., 2011;Araque et al., 2017). As expected from our previous studies (Lafourcade et al., 2007;Lafourcade et al., 2011;Thomazeau et al., 2014), low-frequency tetanic stimulus (10 minutes at 10 hz) induced a long-lasting reduction of the efficacy of excitatory synapses onto layer V/VI pyramidal PFC in wild-type mice. In marked contrast, the same protocol yield to a potentiation of the EPSC in Ts65Dn mice (**Figure 2**).

**Figure 2.**
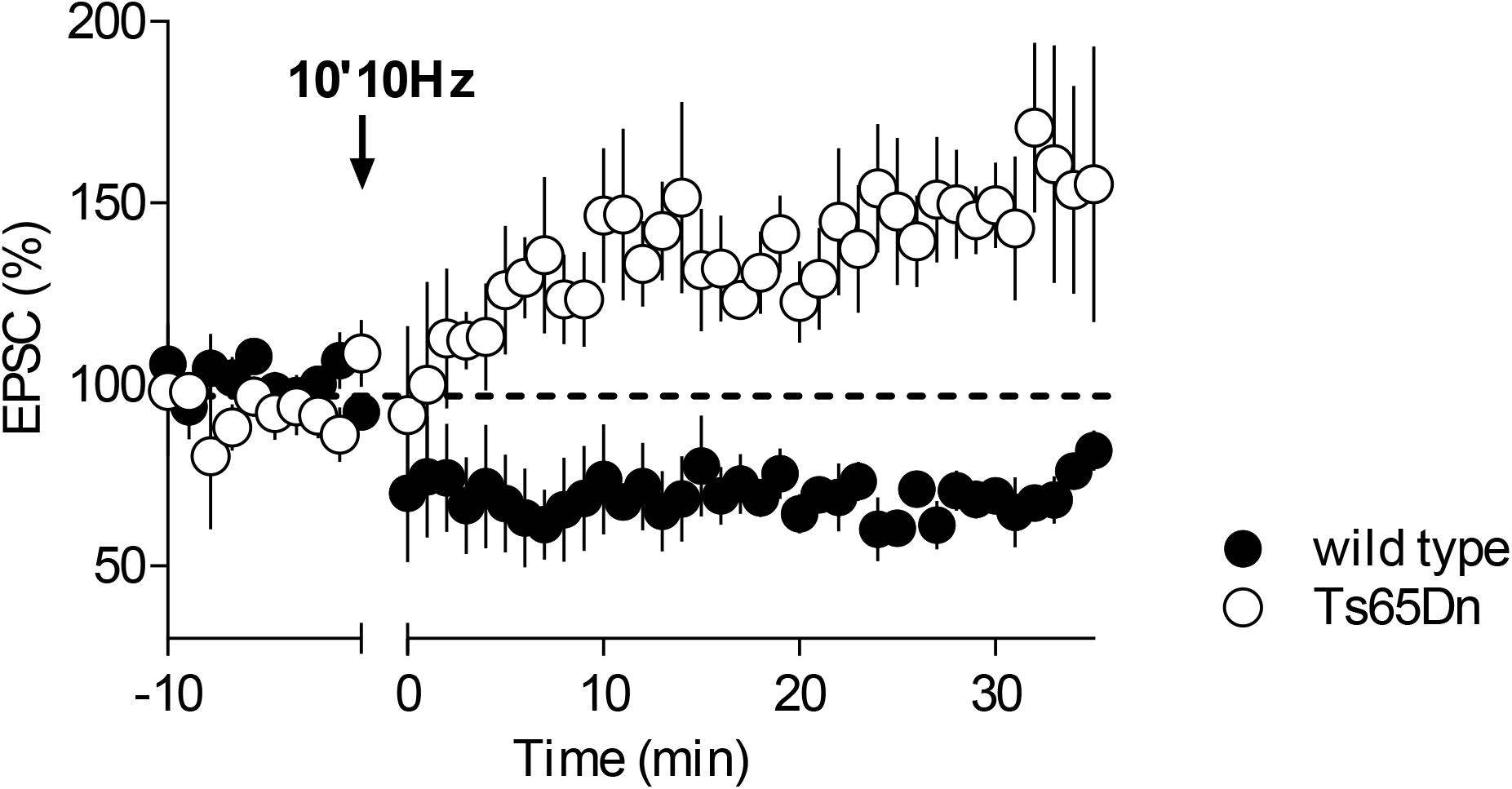
Endocannabinoid-dependent LTD is absent in Ts65Dn mice. Low frequency stimulation of layers II/III induces the long-term depression of excitatory post-synaptic currents recorded in layers V/VI of the PFC in wild-type mice (n = 5, black symbols, 101.28 ± 1.07, baseline versus 72.57 ± 4.08, 30-35 minutes; P = 0.0079, Mann-Whitney test). In Ts65Dn mice this protocol triggered instead a long-term potentiation (n = 5, white symbols, 94.38 ± 2.85, baseline versus 149.47 ± 20.40, 30-35 minutes; P = 0.0079, Mann-Whitney test).

#### 3.2.2. Protein kinase A-mediated long-term potentiation is not changed in Ts65Dn mice

We next examined the presynaptic Adenylyl cyclase (AC)/cAMP-facilitation. Adenylyl cyclase is an enzyme that converts ATP to cAMP, that activates protein kinase A and consequently facilitates transmitter release (Chavez-Noriega and Stevens, 1994;Weisskopf et al., 1994). Synaptic enhancement induced by bath application of the AC activator Forskolin was normal in the aneuploid strain (**Figure 3**), showing that this form of plasticity is not affected in Ts65Dn animals.

**Figure 3.**
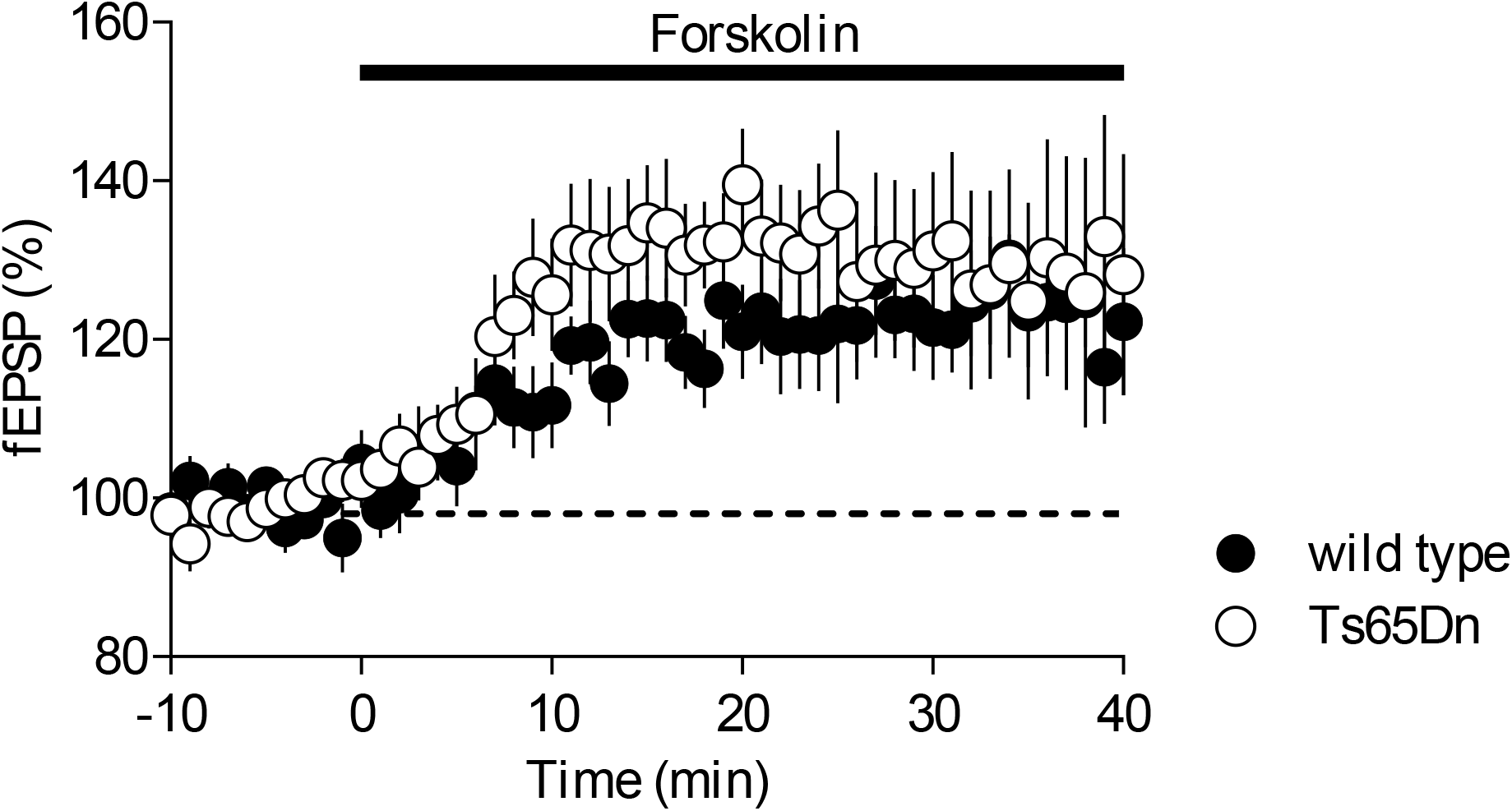
The AC/cAMP-dependent facilitation is intact in Ts65Dn mice. Incubation with the adenylate cyclase activator, forskolin, in the perfusion bath leads to an increase in EPSP recordings in layers V/VI in wild-type mice (n = 8, black symbols, 98.63 ± 0.55, baseline versus 122.75 ± 6.81, 35-40 minutes; P = 0.0104, Mann-Whitney test) and in Ts65Dn mice (n = 8, white symbols, 98.95 ± 0.24, baseline versus 131.97 ± 13.46, 35-40 minutes; P = 0.0205, Mann-Whitney test).

#### 3.2.3. NMDAR-dependent long-term potentiation is intact in Ts65Dn mice

Activity NMDAR-dependent long-term potentiation (LTP) is probably the most widely expressed and extensively studied form of synaptic potentiation (Volianskis et al., 2015). NMDAR-postsynaptic LTP was not altered in Ts65Dn mice (**Figure 4**). Indeed, theta burst stimulation (TBS) of PFC layers II/III induces a similar increase in evoked field potential response of V/VI layers in the two strains of mice.

**Figure 4.**
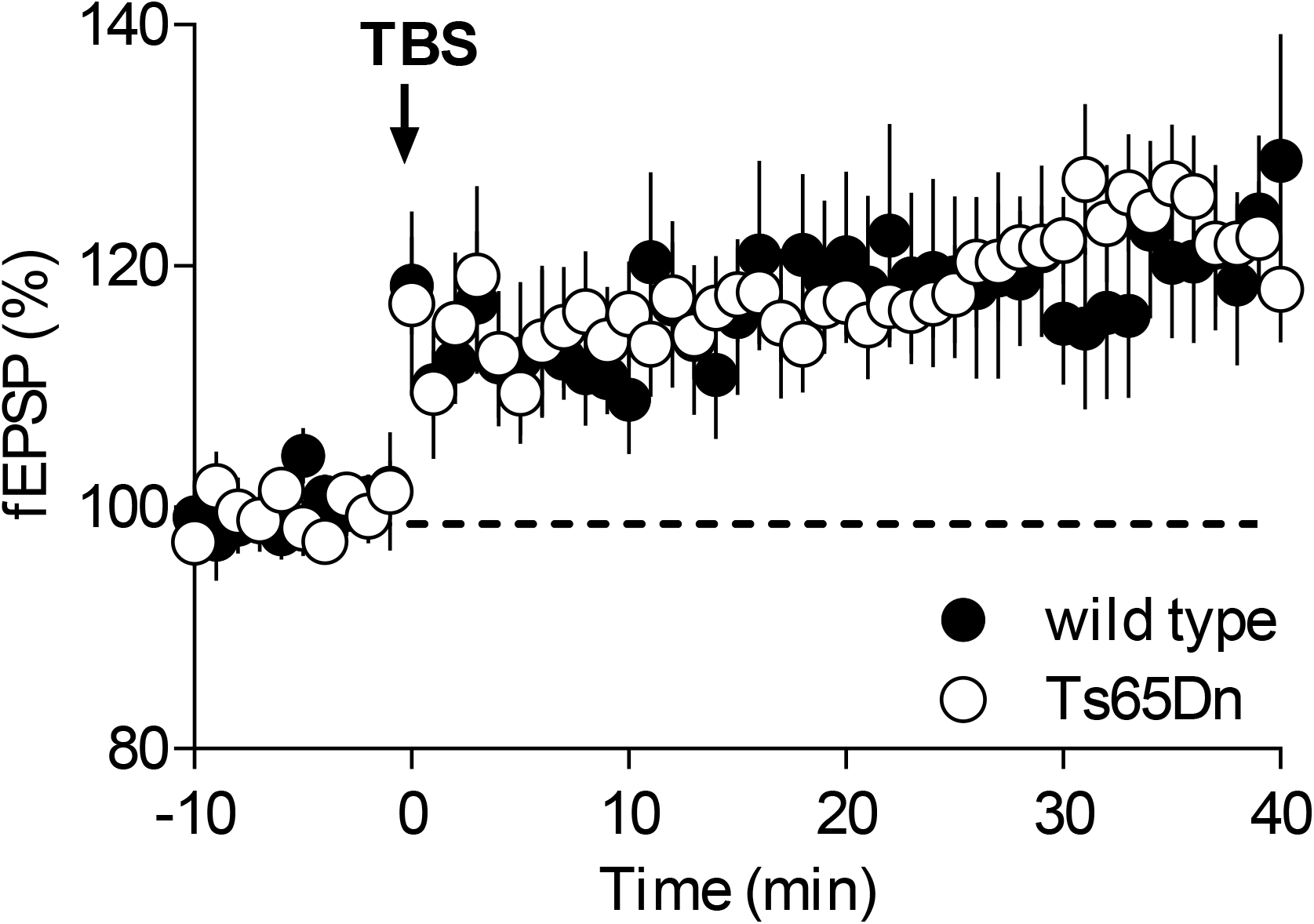
Long-term potentiation dependent on NMDAR is preserved in the Ts65Dn mouse model. A theta burst stimulation of PFC layers II/III induces long-term potentiation of field potentials recorded in layers V/VI in wild-type mice (n = 9, black symbols, 99.82 ± 0.29, baseline versus 119.15 ± 5.88, 35-40 minutes; P = 0.0056, Mann-Whitney test) and in Ts65Dn mice (n = 9, white symbols, 99.53 ± 0.41, baseline versus 123.17 ± 4.49, 35-40 minutes; P = 0.0040, Mann-Whitney test).

## 4. Discussion

In this study, we found altered intrinsic properties and some impaired synaptic plasticity in the PFC of Ts65Dn mice, a model of DS with overexpression of around 130 genes from the mouse analogue of Hsa21, Mmu16, including Dyrk1a. Notably, our results contrast with those reported in mBACtgDyrk1a mice.

Whole-cell patch clamp recordings allowed investigating PFC pyramidal neurons’ cellular intrinsic properties to detect alterations in neuronal excitability. In an earlier study, we showed that the intrinsic properties of PFC layer V/VI pyramidal neurons were not affected in mBACtgDyrk1a mice (Thomazeau et al., 2014). In marked contrast, we report here that Ts65Dn mice exhibit an augmented inward rectification and a higher rheobase (**Figure 1**), reflecting a decrease in neuronal excitability. This is in line with the decreased neuronal excitability already reported in somatosensory cortex layer IV neurons and in dissociated neuronal cultures from the hippocampi of Ts65Dn mice (Cramer et al., 2015;Stern et al., 2015). This reduced intrinsic excitability has been linked to an increase expression level of the inwardly rectifying potassium channel Kir3.2, encoded by the *Kcnj6* gene, one of the triplicated genes in the Ts65Dn mice (Harashima et al., 2006;Best et al., 2007;Best et al., 2012). Strategies to pharmacologically reduce this channel function could be explored to normalize the intrinsic properties of PFC pyramidal neurons in Ts65Dn mice.

We then examined several forms of synaptic plasticity that rely on distinct pre- and postsynaptic mechanisms. Interestingly, eCB-LTD was affected in both mouse models (**Figure 2**; Thomazeau et al., 2014). In contrasts, NMDAR-LTP and AC/cAMP-facilitation were only disrupted in the mBACtgDyr1a mice (**Figures 3 & 4**; Thomazeau et al., 2014; unpublished data). Thus, glutamatergic synaptic plasticity appears to be much less altered in Ts65Dn mice compared to mBACtgDyrk1a mice. Out of all the plasticities tested, only eCB-LTD is impaired. Mechanistically, this form of plasticity is more complex than NMDAR- or mGluR-dependent plasticities, as it involves a series of pre- and postsynaptic loci and processes. This could explain why it is more sensitive to genetic modifications. Surprisingly, the eCB-LTD protocol, lead to LTP in Ts65Dn mice. Various forms of eCB-dependent LTP have been reported in different brain aeras, resulting in an increase neurotransmitter release mediated by either homosynaptic mechanisms, or heterosynaptic processes involving CB1Rs on astrocytes or GABAergic neurons (Navarrete and Araque, 2010;Piette et al., 2020). While endocannabinoids could facilitate the induction of hippocampal LTP by suppressing inhibitory transmission (Carlson et al., 2002), this possibility can be excluded in our experiments, considering that all current experiments were performed in presence of GABA_A_R is pharmacologically blockers.

Numerous studies have shown that LTP is affected in the hippocampus of Ts65Dn mice (Kleschevnikov et al., 2004;Costa and Grybko, 2005;Fernandez et al., 2007;Duchon et al., 2020). The deficit of LTP can be rescued by reducing the magnitude of GABA-mediated signaling through treatment with GABA_A_R antagonist (Kleschevnikov et al., 2004;Costa and Grybko, 2005;Fernandez et al., 2007;Duchon et al., 2020), or by reversing GABA_A_ R signaling, GABA being excitatory rather than inhibitory in Ts65Dn mice (Deidda et al., 2015). In our study, it is possible that the induction of LTP was facilitated by pharmacological inhibition of inhibitory transmission (Thomazeau et al., 2017). It would be compelling to replicate the experiment without the pharmacological blockage of inhibitory transmission to confirm our findings. However, a study conducted on the somatosensory cortex demonstrated that in Ts65Dn mice, the balance between excitatory and inhibitory transmission might remain mostly unaltered, which was indicated by a decrease in the frequency of both excitatory and inhibitory spontaneous synaptic activities (Cramer et al., 2015).

How can we account for the differences in LTP observed between Ts65Dn and mBACtgDyrk1a mice? The partial Ts65Dn model provides a genetic environment more akin to that found in DS, with trisomy of over one hundred genes. However, the regulatory relationships between these genes are currently unknown, and overexpression of genes on chromosome 21 could lead to modified expression of genes on other chromosomes. While comparative analyses of gene expression on other chromosomes have demonstrated that gene deregulations are specific to Hsa21 in DS foetuses (Mao et al., 2003), research on the Ts65Dn model has shown that there is a global alteration in gene expression in these mice (Chrast et al., 2000;Lyle et al., 2004). Furthermore, gene products can interact with one another, creating compensatory or exacerbating mechanisms, which can affect synaptic parameters in Ts65Dn mice differently, reflecting the intricacy of the gene effects associated with this condition. In simpler terms, the deficits observed in Ts65Dn mice that are not present in mBACDyrk1a mice could be attributed to detrimental mechanisms arising from the overexpression of genes other than Dyrk1a. Conversely, the absence of defects in Ts65Dn that are seen in mBACDyrk1a mice could indicate the synergistic and/or compensatory activities of these three-copy genes. According to Dowjat et al. (Dowjat et al., 2007), the amount of DYRK1A protein in the brains of Ts65Dn animals is 1.5 times higher. However, it is important to confirm whether this rate is consistent specifically within the PFC of the Ts65Dn animals. Currently, little is known about the spatiotemporal regulation of DYRK1A expression, or whether the overexpression of Dyrk1a and other trisomal genes in mouse models of DS varies across age, sex, and brain region throughout development (Stringer et al., 2017). Recent research has shown that the expression of DYRK1A protein in P15 mice is sex-dependent in Ts65Dn mice, with significant overexpression in males but not in females (Hawley et al., 2022). This suggests that brain development in male and female Ts65Dn mice may have different trajectories, underscoring the significance of conducting future DS mouse model studies that include both males and females.

It is important to note that the Ts65Dn animals have triplication of 50 non-orthologous Mmu17 genes in addition to Mmu16 orthologous genes, which may complicate the genotype/phenotype relationship and contribute to the observed synaptic phenotypes. To address this issue, future experiments could be performed on the refined Ts65Dn model recently developed, the Ts66Yah, which retains the major features of DS, but showed an overall milder phenotype than Ts65Dn mice (Duchon et al., 2022). However, it is worth noting that the partial Ts65Dn model does not overexpress the Hsa21 orthologous genes carried by Mmu10 and Mmu17, which raises questions about the relevance of this partial mouse model as a standard for DS research (Guedj et al., 2022). Therefore, it is important to conduct comparative studies in different mouse models for DS, each with its limitations in recapitulating the full spectrum of human DS. To fully compare the Ts65Dn and mBACtgDyrk1a mouse models, additional experiments such as examining basal glutamatergic transmission, mGluR3-LTD, dendrites and spines morphology would be necessary (see Thomazeau et al., 2014).

Overall, the parallel between this more complete Ts65Dn DS model and the monogenic Dyrk1a overexpression model highlights the complexity of the physiological and pathological mechanisms responsible for the neurocognitive symptoms of DS.

## 6. Acknowledgements

We thank members of the Manzoni lab for help and useful discussions, and the National Institute of Mental Health’s Chemical Synthesis and Drug Supply Program (Rockville, MD, USA) for providing DNQX.

## 9. Author Contribution

A.T. and O.J.M. designed the research. AT and OL carried out the experiments. A.T. analyzed the data. AT and OJM wrote the manuscript.

## 10. Data Availability Statement

The raw data supporting the conclusions of this manuscript will be made available by the authors, without undue reservation, to any qualified researcher.

